# Protein-level mapping reveals novel Immune State Types (ISTs) associated with *Chlamydia trachomatis* infection, inflammatory remodeling, and scarring in trachoma-hyperendemic Amhara Ethiopia

**DOI:** 10.64898/2026.09.30.755767

**Authors:** Xiaoyi Charlotte Shi, Olusola Olagoke, Brianna Renatte Titiheruw, Seongwon Chung, Hiwot Degineh Mengistie, Kaleb Asfaha, Timothy D. Read, Deborah Dean

## Abstract

**Background:** Trachoma remains the leading infectious cause of blindness. Over 100M people are at risk, and 1.9M already live with visual impairment or blindness. Ethiopia bears the greatest burden of disease with ∼61M or 59% of the known affected global population. Although *Chlamydia trachomatis* (*Ct*) infection initiates disease, infection alone fails to explain why inflammation resolves in some but progresses to irreversible scarring and vision loss in others. Previous studies have largely examined gene expression or limited protein panels, leaving the conjunctival immune protein environment poorly characterized. We hypothesized that distinct conjunctival immune states capture these pathogenic processes and could reveal biomarkers for selecting treatment and surveillance strategies.

**Methods and Findings:** We conducted a cross-sectional study of 667 participants residing in the hyperendemic Amhara region of Ethiopia at least one year after the most recent azithromycin mass drug administration to treat *Ct*. The upper tarsal conjunctivae was graded for TF and TI, trachomatous inflammation-follicular or intense, respectively; I-TS/TT, inflammation with scarring with or without trichiasis, defined as one or more eyelashes touching the globe of the eye; and TS and TT, scarring with or without trichiasis but no inflammation. T0 represented no trachoma. The protein concentrations of 41 immune mediators were quantified using Meso Scale Discovery electrochemiluminescence immunoassays. Associations with infection and grade were evaluated using covariate-adjusted models with false discovery rate correction. Bernoulli mixture modeling (BMM) revealed four distinct Immune State Types (ISTs). Both approaches supported a shared biological framework characterized by selective immune activation in *Ct* infection, broad inflammation in TF/TI, and chemokine remodeling versus immune quiescence in the scarred conjunctiva. *Ct* infection was associated with a focused IFN-γ-centered response. IST1 captured this immune response where 51.9% of the participants had *Ct*; we therefore termed IST1 “Quiescent Infection.” In contrast, TF/TI was associated with higher concentrations of proinflammatory, Th1/Th17, chemokine, regulatory, and remodeling pathways mediators. This profile fit well within IST2, termed “Broad Inflammation”, where 72.4% of the participants had TF/TI. TS/TT retained considerable chemokine activity, whereas I-TS/TT showed a narrower, IL-17A-dominant inflammatory profile. BMM separated scarring into two states characterized by distinct profiles: IST3, “Conjunctival Remodeling,” with significantly stronger chemokine and T-cell maintenance signals; and IST4, “Quiescent Scarring,” with the majority of chemokines being significantly downregulated. This suggests that scarred conjunctiva can retain substantial immune activity or shift toward a more attenuated state.

**Conclusions:** This study establishes the first IST framework for trachoma, revealing biologically distinct immune states with different potential clinical implications. IST1 indicates the need for antibiotic treatment, whereas IST2 suggests that novel immunotherapeutics might be effective in thwarting disease progression. The upregulated chemokines in IST3 may indicate risk of progression to TT necessitating prospective studies and potential novel antifibrotic interventions. Together, our findings provide a strong biological foundation for stratifying patients by immune state and matching them to preventive and therapeutic care. Prospective validation and streamlined biomarker panels will support the translation of this framework into field settings to help avert disease progression and blindness.

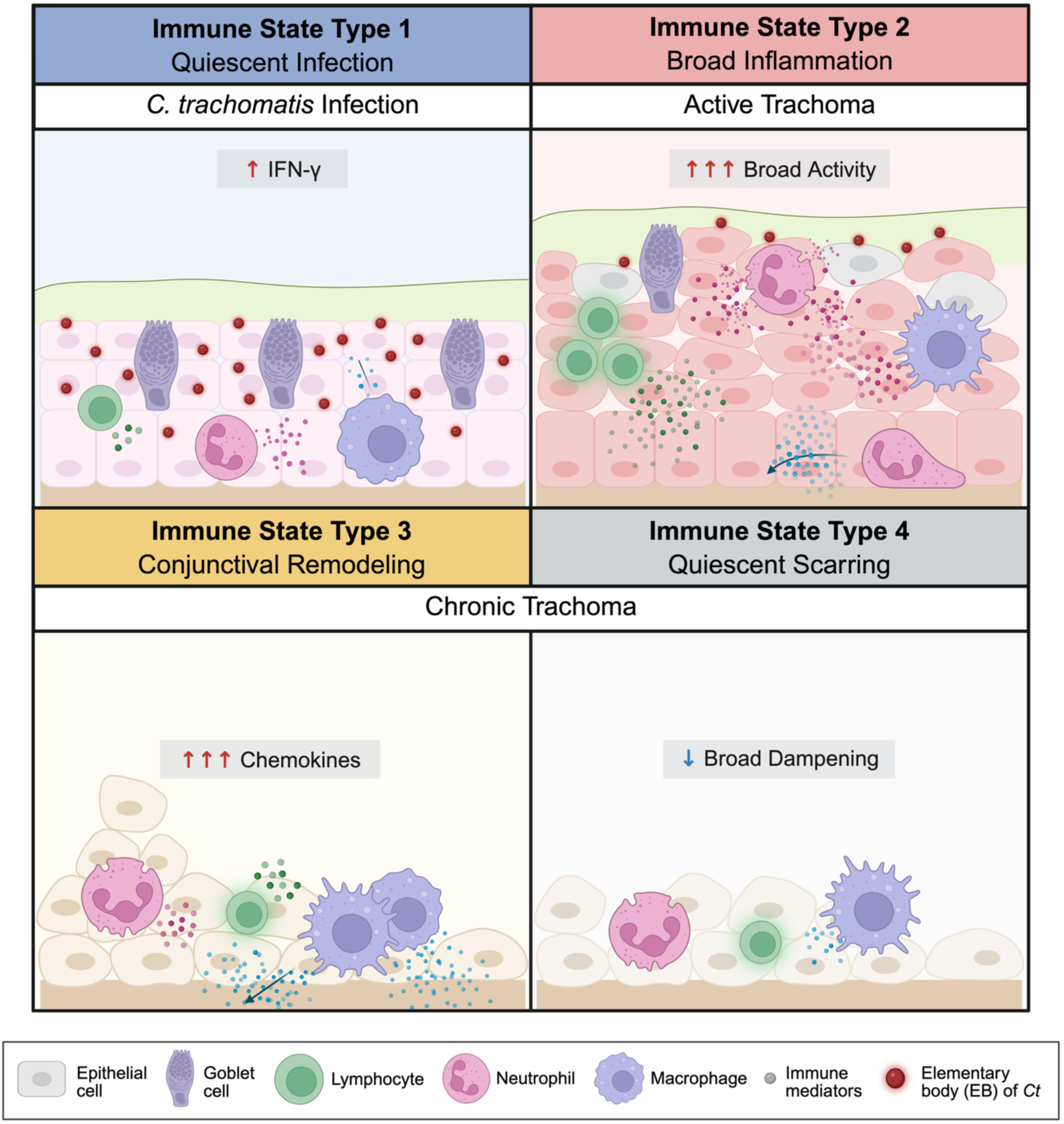
GRAPHICAL ABSTRACT.

## INTRODUCTION

Trachoma is a chronic ocular disease caused by the obligate intracellular bacterium *Chlamydia trachomatis* (*Ct*) and remains the leading cause of infectious blindness worldwide [1]. Infection in childhood may be mild or clinically silent, yet repeated episodes can drive inflammation and subsequent conjunctival scarring [2]. Scarring can lead to inturned eyelashes— referred to as trichiasis—that abrade the cornea and heal by fibrosis causing visual impairment or blindness [2]. Despite decades of global control efforts under the World Health Organization (WHO)-endorsed SAFE (**S**urgery for blinding trachoma, **A**ntibiotics to clear infection, **F**acial cleanliness, and **E**nvironmental improvement) strategy, trachoma remains a public health concern in over 30 countries, causing blindness or visual impairment in ∼1.9 million people [1]. Approximately 100M people continue to live in trachoma-endemic areas and remain at risk of vision loss, though this figure is likely much higher due to structural surveillance gaps [1].

A central unresolved question in trachoma pathogenesis is why some individuals clear infection without lasting tissue damage, whereas others develop inflammation, progressive scarring, and blinding disease. Although *Ct* initiates disease, infection alone does not explain disease progression. Vision-threatening trachoma likely reflects host immune responses to repeated infection-induced inflammation with subsequent epithelial dysfunction and tissue remodeling [3]. Defining protective versus pathogenic responses is therefore essential for understanding trachoma pathogenesis and developing vaccines, control strategies, and novel therapies.

The majority of previous studies of trachoma immunology have relied on mRNA expression studies, which suggested that active *Ct* infection was associated with a T helper 1 (Th1) cell-mediated inflammatory response, including increased interferon (IFN)-γ and innate immune mediators interleukin (IL)-1β and tumor necrosis factor (TNF) [4–11]. Because inflammation can persist after infection can no longer be detected, the data supported a model in which trachoma evolves from infection-driven inflammation toward partially infection-independent scarring.

However, transcript-level profiling provides an incomplete view of the immune environment because mRNA abundance is shaped by post-transcriptional regulation, translational control, secretion dynamics, protein stability, and local tissue context and does not reliably predict protein concentrations [12]. Proteogenomic studies have consistently shown only modest concordance between mRNA expression and protein concentration with mean Spearman correlations ranging from 0.23 to 0.45 [13–15]. Protein-level measurements are therefore needed to define the immune signals that are actually present in the conjunctivae.

Prior studies of immune proteins in trachoma populations have used narrow panels of 10 to 25 analytes and have not examined key cytokine families like the IL-17 axis (IL-17A-F), which contributes to mucosal barrier immunity, neutrophil recruitment, and inflammatory tissue remodeling [16]. In Nepal, Luminex-based protein profiling found no IFN-γ differences by infection or inflammation, but conjunctival scarring was associated with a chemokine-enriched profile, including macrophage inflammatory protein (MIP)-1α, MIP-1β, monocyte chemoattractant protein (MCP)-1, and eotaxin [17]. In Tanzania, scarring was associated with higher tear IL-8, CXCL1, IL-10, and lysozyme levels, but these associations were less clearly defined in The Gambia [18], suggesting that protein-level profiles may reveal immune responses that vary across populations.

Ethiopia has the highest known trachoma burden, affecting an estimated 61 million people, or 59% of the known global total [19]. Despite ∼15 years of SAFE implementation, including azithromycin mass drug administration (MDA) and environmental interventions, hyperendemic regions remain [20], suggesting a gap in our understanding of infection and host immune responses that sustain inflammation and promote scarring in elimination-resistant settings. Since few studies have examined the immune response associated with trachoma in this region, and only narrowly on trachoma-related scarring [21,22], broader immunologic research is needed.

Here, we used a highly sensitive electrochemiluminescence immunoassay (i.e., Meso Scale Discovery (MSD)) to quantify 41 immune mediator proteins in conjunctival samples from 667 villagers in the trachoma-hyperendemic Amhara region of Ethiopia. This expanded panel captures proinflammatory, anti-inflammatory, T helper 1/T helper 2 (Th1/Th2), T helper 17 (Th17), T-cell regulatory, chemokine, and growth factor proteins. We examined immune analytes using unsupervised mixture modeling to identify novel ocular Immune State Types (ISTs) associated with population characteristics including infection, inflammation, and scarring, providing a robust framework for understanding trachoma pathogenesis.

## METHODS

### Study Population, Trachoma Grading, Ocular Sample Collection, and *Ct* Detection

The study was conducted across three village regions in Amhara, Ethiopia, at least one year following the most recent MDA program as we have described [23]. Individuals aged one year or older were consented prior to enrollment as per the Institutional Review Boards of the University of California, San Francisco School of Medicine, and the Ethiopian Ministry of Health.

The metadata for this study are shown in Supplemental **Table S1**. High-resolution photos of the upper tarsal conjunctiva of both eyes were taken before conjunctival swabbing to confirm the trachoma grading performed in the field as previously described [24]. Briefly, using a 2.5x ocular loupe, trained ophthalmic personnel examined the upper tarsal conjunctivae and assigned trachoma grades using the amended WHO grading scale [25]: trachomatous inflammation-follicular (TF), ≥5 follicles of at least 0.5-mm; trachomatous inflammation-intense (TI), pronounced inflammation obscuring over half of the underlying vessels; trachomatous scarring (TS), white visible scars; and trachomatous trichiasis (TT), ≥1 eyelash touching the eyeball or eyelash epilation. Participants were classified into four mutually exclusive clinical categories: T0, indicating no evidence of trachoma; TF/TI, indicating TF or TI without TS or TT; TS/TT, indicating TS with or without TT without TF or TI; and I-TS/TT, indicating TF or TI occurring concurrently with TS with or without TT.

Conjunctival samples were obtained from the upper tarsus of each eye using FLOQ swabs (Copan, Murrieta, CA) as we described [24]. Swabs were placed into sterile M4 medium (Remel, Lenexa, Kansas) and stored at −80°C until processed in the MSD assay as previously described [26].

*Ct* infection status was determined using a two-step approach. A real time (RT)-quantitative (q)PCR assay was used to detect *Ct* as described previously [27]. Metagenomic shotgun sequencing data generated as part of the parent study [23] were also used to detect *Ct* genomic reads [23,28]. Briefly, *Ct* positivity was defined as ≥1 genome copy by qPCR and/or >1 sequencing read identified by Kraken2 or CZ ID and validated by Megablast with 99-100% identity to the *Ct* genome or plasmid.

Participant occupations were collected in the field through surveys and classified using the same categories as our recent study of the same cohort: Children, Student, Cooking/Childcare, Compound Duties, Agriculture, Service, Trade & Manual Labor, Professional/Technical, and Other [28].

### Multiplex Immunoassay Measurement of Immune Analytes in Conjunctival Samples

Human Cytokine/Chemokine V-PLEX kits (MSD, Rockville, MD) were used to quantify the analytes listed in **Table 1** as we described previously [26]. Briefly, 150 µL of M4 transport medium per sample was diluted 1:2 or 1:4 prior to adding to the assay plate; serial four-fold dilutions of seven standards with a final standard of diluent alone were added to separate wells. Plates were processed with antibody and washed prior to adding the SULFO-TAG detection antibody cocktail. MSD GOLD Read Buffer B was dispensed into each well after washing, and signal acquisition was performed using the MESO QuickPlex SQ 120MM platform (MSD, Rockville, MD).

**Table 1.**
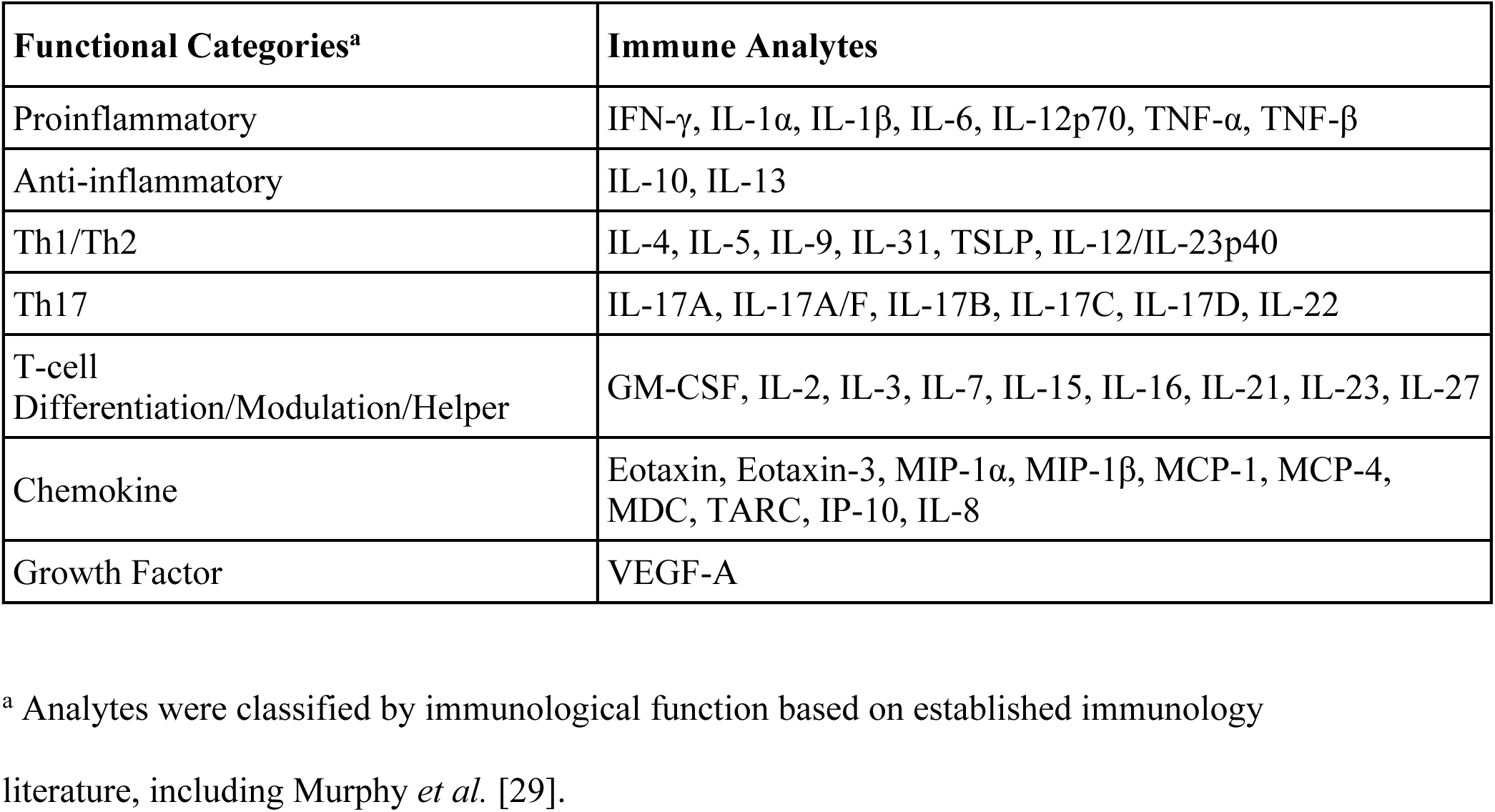
Immune analyte panels measured in this study and corresponding immune function.

| **Functional Categories<sup>a</sup>** | **Immune Analytes** |
| --- | --- |
| Proinflammatory | IFN-γ, IL-1α, IL-1β, IL-6, IL-12p70, TNF-α, TNF-β |
| Anti-inflammatory | IL-10, IL-13 |
| Th1/Th2 | IL-4, IL-5, IL-9, IL-31, TSLP, IL-12/IL-23p40 |
| Th17 | IL-17A, IL-17A/F, IL-17B, IL-17C, IL-17D, IL-22 |
| T-cell<br>Differentiation/Modulation/Helper | GM-CSF, IL-2, IL-3, IL-7, IL-15, IL-16, IL-21, IL-23, IL-27 |
| Chemokine | Eotaxin, Eotaxin-3, MIP-1α, MIP-1β, MCP-1, MCP-4, MDC, TARC, IP-10, IL-8 |
| Growth Factor | VEGF-A |

<sup>a</sup> Analytes were classified by immunological function based on established immunology literature, including Murphy *et al.* [29].

For each assay plate, the standards were used to generate the corresponding standard curve to convert raw signals from experimental wells into concentrations reported in pg/mL using the Discovery Workbench 4.0 analysis software (MSD, Rockville, MD). Readings below the assay’s lower limit of detection were treated as not detectable and recorded as 0. Readings above the upper limit of detection were given the highest concentration that remained within the measurable range of the assay.

### Analysis of Immune Protein Associations with Trachoma Grade, *Ct* Infection, Age Group and Sex

To compare conjunctival immune analyte concentrations across clinical and infection groups while accounting for potential confounders, associations with trachoma grade and with *Ct* infection status were evaluated using analysis of covariance (ANCOVA) in R [30]. Concentrations were transformed as log_2_ (concentration + 1) before analysis to normalize the data. Model estimates represent adjusted differences in log_2_-transformed analyte concentrations between comparison groups and included age, sex, and either *Ct* infection status or trachoma grade as covariates, depending on the comparison. Age was included as a categorical covariate and classified into four groups: children (<10 years), adolescents (10–19 years), adults (20–49 years), and older adults (>49 years). For each comparison, *P* values were adjusted for multiple testing across 41 analytes using the Benjamini-Hochberg (BH) procedure to control for false discovery rate (FDR) [31].

To assess the effect of *Ct* infection on the host immune response, analyte concentrations were compared for *Ct* infection versus no infection independently for each trachoma grade (i.e., T0, TF/TI, I-TS/TT, and TS/TT). For each analyte, samples within a specific grade were ranked by concentration, and Spearman’s correlation test was used to determine whether *Ct*-positive samples had enriched or depleted ranks compared with uninfected samples. *P* values were adjusted for FDR using the BH procedure.

### Bernoulli Mixture Modeling of Immune Analyte Detection Patterns

Bernoulli mixture modeling (BMM) was used to identify latent patterns of immune analyte detection across samples [32]. This probabilistic clustering approach is designed for binary data and assumes that samples arise from one of *K* latent classes. Because cluster membership is unknown, the model estimates each sample’s probability of belonging to each cluster. These probabilities are used to assign samples to the most likely cluster and to estimate the relative size of each cluster. Within each cluster, the model also estimates the probability that each analyte is detected, providing a summary of the characteristic immune detection pattern for that cluster [32]. To assess robustness to modelling assumptions, we compared the Bernoulli mixture with several alternative approaches that relaxed or avoided its key assumptions: a mixture of Rasch models accounting for sample-specific detection propensity [33]; non-negative matrix factorization of the concentration matrix, with samples grouped according to their dominant latent factor [34]; compositional analysis of concentrations using a centred log-ratio transformation followed by Gaussian mixture modelling [35,36]; a left-censored (Tobit) mixture model retaining measured magnitudes [37,38]; and hierarchical clustering based on Jaccard and Jensen-Shannon distances [39].

Immune analyte concentrations were converted to binary values coded as 1 for detected and 0 for not detected. BMM models were fitted in R using the “flexmix” package [40–43]. Models with K = 2–8 components were evaluated, and the optimal model was selected using the Bayesian information criterion (BIC) [40]. To reduce sensitivity to initialization, 50 random starts were used for each value of K, and expectation-maximization iterations were allowed to run for 1000 iterations. Cluster assignments were obtained from the selected model. Cluster-level detection profiles were summarized by the mean binary value for each analyte within each cluster, corresponding to the detection frequency of that analyte.

Sample detection breadth was defined as the number of the 41 analytes detected in each sample, whereas analyte prevalence was defined as the proportion of samples in which each analyte was detected. To determine whether the data contained a nonrandom clustered structure, 50 null matrices were generated that preserved sample detection breadth and analyte prevalence while randomizing analyte co-occurrence within each sample. Bernoulli mixture models were fitted to the observed and null matrices, and their BIC improvements over K = 1, the one-cluster model, were compared.

To determine whether IST separation was driven mainly by sample detection breadth or by analyte identity, breadth was compared across ISTs using a Kruskal-Wallis test followed by unadjusted two-sided pairwise Wilcoxon rank-sum tests. Multinomial logistic regression predicted IST from either breadth alone or all 41 binary detection indicators. Samples were randomly divided into five approximately equal folds. Using the same five random folds, each model was trained on four folds and tested on the fifth, with accuracy defined as the proportion correctly classified.

Following these post-hoc assessments of IST structure, age distributions across ISTs were compared using the Kruskal-Wallis test, followed by pairwise Wilcoxon rank-sum tests with BH correction. Enrichment or depletion of trachoma grade, *Ct* infection status, occupation category, and village region within each IST was evaluated using 2 × 2 IST-versus-rest contingency tables. Chi-square tests were used when expected cell counts were sufficient (≥ 5), and two-sided Fisher’s exact tests were used for small, expected counts (<5). Risk ratios and 95% confidence intervals (CI) were calculated using the Katz log method.

## RESULTS

### Study population, trachoma grading, and detection of *Ct*

Metadata for the participants in this study are shown in **Supplementary Table S1**. Participant characteristics are summarized in **Table 2**. The cohort of 667 participants was enriched for chronic trachoma with most participants classified as TS/TT (70.31%; 469/667) or I-TS/TT (5.70%; 38/667). The age distribution followed the expected natural history of trachoma: TF/TI participants were the youngest (median age 4.5 years), whereas TS/TT and I-TS/TT participants were the oldest (median age 55 and 50 years, respectively). TF/TI participants were significantly younger than T0 participants, while TS/TT and I-TS/TT participants were significantly older than both T0 and TF/TI participants, consistent with active inflammatory disease occurring primarily in children, and scarring disease accumulating later in life [44]. Females had a significantly higher risk of I-TS/TT compared with males. Overall, 231 (34.6%) of 667 participants were *Ct*-positive; infection was frequent among TF/TI participants (53.45%; 31/58) and significantly higher in TF/TI than in T0 participants and compared to TS/TT.

**Table 2.** Characteristics of the study population by age, sex, trachoma grade and *Ct* infection.

| **Trachoma Grade** | **T0**<br>**(n=100)** | **TF/TI**<br>**(n=58)** | **I-TS/TT**<br>**(n=38)** | **TS/TT**<br>**(n=469)** | **Missing**<br>**grade (n=2)** | **Total**<br>**(n=667)** |
| --- | --- | --- | --- | --- | --- | --- |
| **Age in years<sup>a</sup>**<br>**[Median (range)]** | 20 (1-70) | 4.5 (1-60) | 50 (17-80) | 55 (6-90) | 58.5 (53-64) | 48 (1-90) |
| **Male** | 62 | 33 | 12 | 250 | 2 | 359 |
| **Female<sup>b</sup>** | 38 | 25 | 26 | 219 | 0 | 308 |
| ***Ct* infection<sup>c</sup>** | 32/100<br>(32.0%) | 31/58<br>(53.4%) | 15/38<br>(39.5%) | 152/469<br>(32.4%) | 1/2 | 231/667<br>(34.6%) |

<sup>a</sup> Individuals with TF/TI were significantly younger than those with T0 (FDR-adjusted $P = 2.31 \times 10^{-5}$ ), while those with TS/TT or I-TS/TT were significantly older than those with T0 (FDR-adjusted $P = 9.30 \times 10^{-44}$ and $5.91 \times 10^{-13}$ ) and TF/TI (FDR-adjusted $P = 5.21 \times 10^{-32}$ and $2.97 \times 10^{-18}$ ), respectively, by Welch's t-test.
<sup>b</sup> Females had a significantly higher risk of I-TS/TT compared with males (risk ratio (RR), 2.51; 95% confidence interval (CI), 1.29-4.89; FDR-adjusted $P = 0.0196$ ), by chi-square test.
<sup>c</sup> TF/TI was associated with higher risk of *Ct* compared with T0 (RR, 1.67; 95% CI 1.15-2.43, FDR-adjusted $P = 0.0239$ ) and with TS/TT (RR, 1.65; 95% CI: 1.25-2.17; FDR-adjusted $P = 0.00899$ ), by chi-square tests.

### *Ct* infection was associated primarily with a Th1/Th2 response

*Ct* infection was associated with significantly higher concentrations of IFN-γ, IL-13, IL-9, thymic stromal lymphopoietin (TSLP), and IL-17B compared with no infection after adjustment for age, sex, and trachoma grade (**Fig 1**). TSLP is an epithelial mediator of Th2 responses, while IL-13 and IL-9 are Th2 cytokines [45]. IFN-γ is a signature proinflammatory Th1 cytokine that mediates immunity against intracellular bacteria and is consistently found to be upregulated during *Ct* infection [4–11].

**Fig 1.**
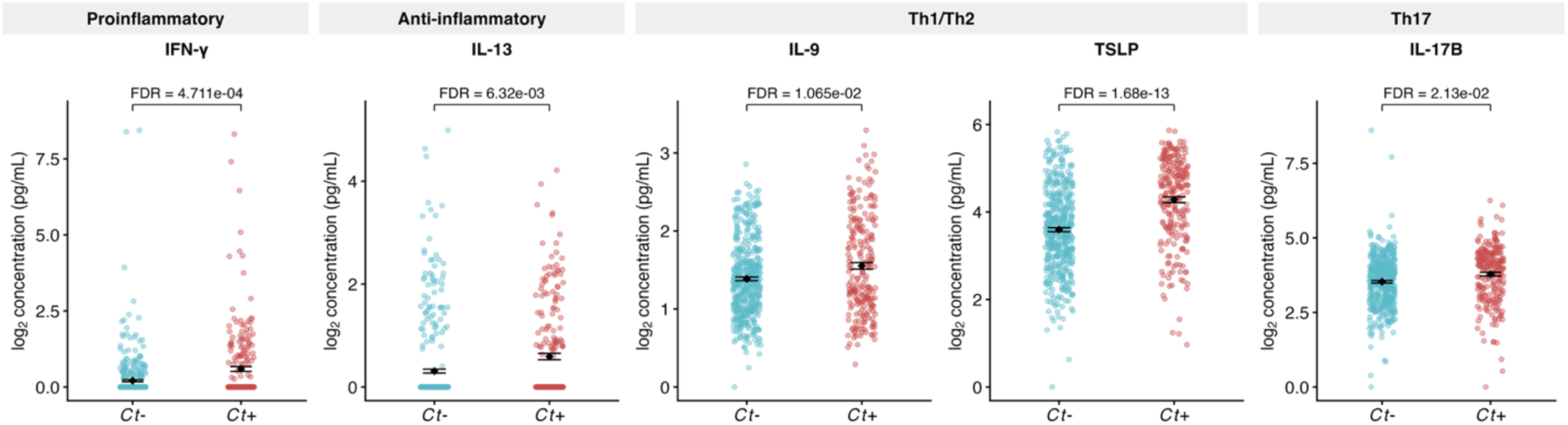
Elevation of Th1/Th2-associated immune mediators in *Ct* infection. Cytokine concentrations are shown for *Ct*-negative (n=434) and *Ct*-positive samples (n=230) after log_2_ transformation. Samples without an age (n=1) or trachoma grade (n=2) were excluded. Each point represents an individual sample. Group mean and standard error of the mean are shown. Differences between *Ct*-negative and *Ct*-positive samples were assessed using covariate linear models adjusting for age, sex, and trachoma grade. FDR-adjusted *P* values are shown above each comparison.

### TF/TI participants had broad inflammatory and Th1/Th17-associated mucosal immune activation, whereas I-TS/TT participants had a narrower Th17-associated immune profile

Compared with T0, participants with TF/TI had significantly higher concentrations of Th1-associated cytokines IFN-γ and TNF-β and other proinflammatory cytokines, including IL-6, IL-1α, IL-1β, and TNF-α after adjustment for age, sex, and *Ct* infection status (**Fig 2**), consistent with prior trachoma studies showing increased inflammatory cytokine mRNA expression [4–7], immunoreactivity [46], or protein levels [17] during active disease. TF/TI was also significantly associated with higher concentrations of Th17-axis-related mediators, including IL-17A, IL-17B, IL-17C, IL-17D, and IL-22, in addition to the regulatory and T-cell modulatory analytes IL-16, IL-15, granulocyte-macrophage colony-stimulating factor (GM-CSF), IL-12/IL-23p40, IL-10, and the growth factor vascular endothelial growth factor A (VEGF-A). Most measured chemokines, including eotaxin, IL-8, monocyte chemoattractant protein (MCP)-1, macrophage inflammatory protein (MIP)-1α, and MIP-1β, were significantly elevated. Macrophage-derived chemokine (MDC), interferon gamma-induced protein (IP)-10, and eotaxin-3 showed borderline increases (FDR = 0.08, 0.095, 0.0996, respectively), indicating a broad response involved in tissue remodeling and fibrosis.

**Fig 2.**
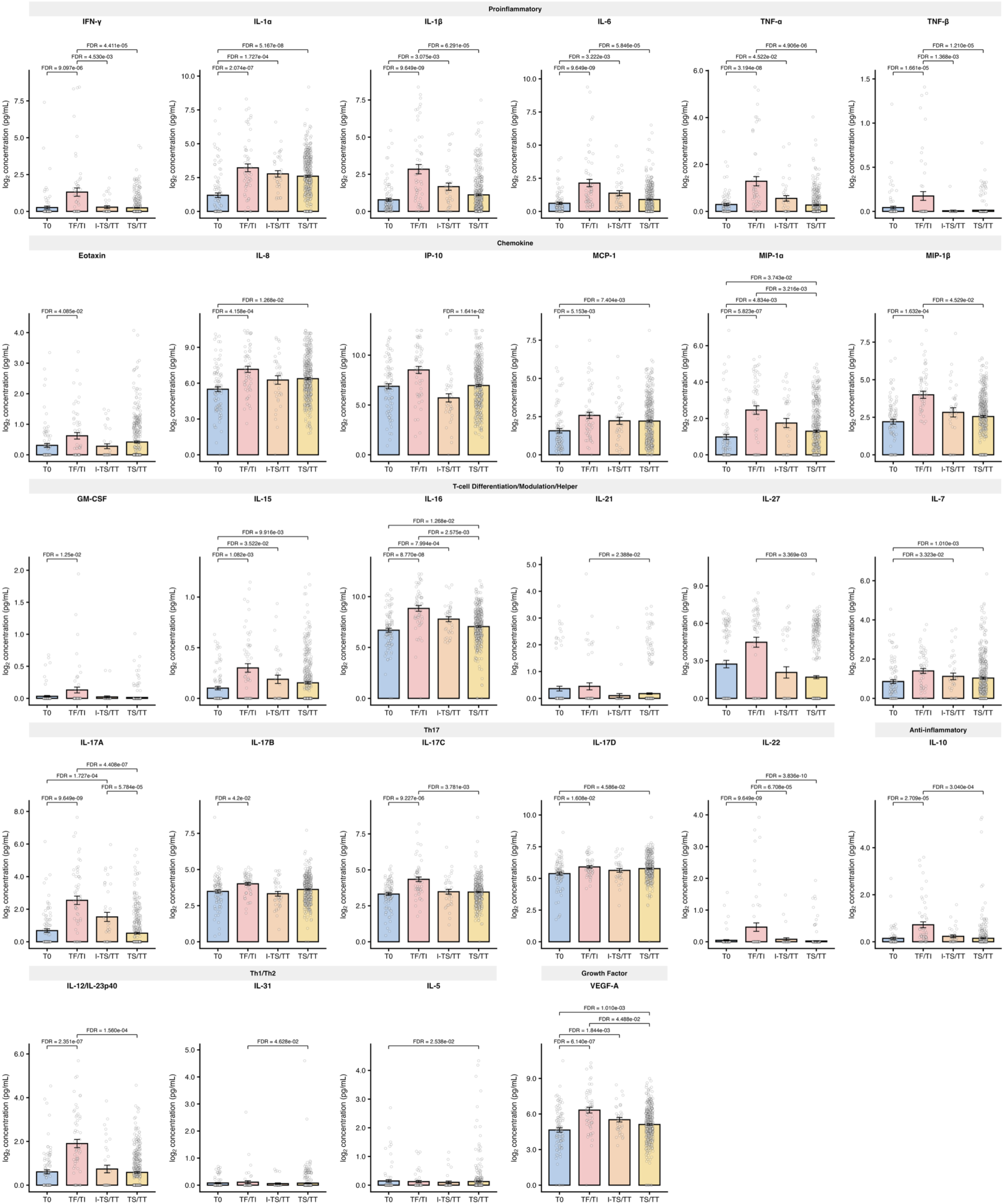
Immune analyte concentrations differ across clinical trachoma grades. Log_2_-transformed analyte concentrations are shown for T0 (n=99), TF/TI (n=58), I-TS/TT (n=38), and TS/TT (n=469). Samples without an age (n=1) or grade (n=2) were excluded. Each point represents an individual sample. Colored bars indicate group means, and error bars represent the standard error of the mean. Pairwise differences between grades were assessed using covariate-adjusted models controlling for age group, sex, and *Ct* infection status. Brackets indicate significant pairwise differences, with FDR-adjusted *P* values shown above each comparison. Overall, immune analyte concentrations were generally lowest in T0 and highest in TF/TI, while TS/TT showed attenuated immune activation and I-TS/TT had an intermediate but distinct profile.

By contrast, I-TS/TT participants had a narrower profile with Th17, but not Th1, features **(Fig 2)**. Although a similar group of general proinflammatory cytokines were elevated (IL-1α, IL-1β, IL-6, and TNF-α), the Th1-associated cytokines IFN-γ and TNF-β were not. Instead, I-TS/TT was associated with elevated IL-17A, the T-cell modulators IL-7, IL-15, and IL-16, VEGF-A, and the chemokine MIP-1α, with a trend towards significant MIP-1β and MCP-1 levels (FDR = 0.07).

### TS/TT was associated with inflammation and chemokine enrichment relative to T0

Compared with T0, TS/TT was associated with significantly higher concentrations of a smaller set of immune analytes after adjusting for age, sex, and *Ct* infection status (**Fig 2**). These included the proinflammatory cytokine IL-1α; the Th2 cytokine IL-5; T-cell modulatory cytokines IL-7, IL-15, and IL-16; chemokines IL-8, MCP-1, MIP-1α; and growth factor VEGF-A as well as IL-17D. The chemokines eotaxin, eotaxin-3, and MIP-1β were also elevated but not quite significant (FDR = 0.07). Together, these immune mediators suggest persistent inflammation, ongoing tissue remodeling, and fibrogenesis that occurs during scarring.

### TS/TT showed attenuation of TF/TI-enriched inflammatory signals, while I-TS/TT had a distinct intermediate profile

TS/TT was compared to TF/TI to distinguish the immune profiles of active and chronic trachoma. All analytes that differed were significantly lower in TS/TT, and most were also elevated in TF/TI relative to T0, except for IL21, IL27, and IL31 (**Fig 2**). These included the proinflammatory mediators IFN-γ, TNF-β, TNF-α, IL-6, and IL-1β; Th17-associated cytokines IL-17A, IL-17C, and IL-22; and T-cell-related mediators IL-12/IL-23p40, IL-16, IL-21, IL-27, and IL-31. TS/TT also had lower IL-10, suggesting weaker anti-inflammatory regulatory counter-signaling, as well as chemokines MIP-1α and MIP-1β and the growth factor VEGF-A. Together, these findings indicate broadly attenuated immune activation in TS/TT compared with TF/TI.

I-TS/TT had an intermediate profile distinct from both TF/TI and TS/TT. Compared with TF/TI, I-TS/TT had lower concentrations of the Th1 cytokines IFN-γ and TNF-β and the Th17-associated cytokine IL-22, indicating reduced Th1 and some Th17-associated signaling. Compared with TS/TT, however, I-TS/TT had higher IL-17A but lower IP-10 concentrations. Thus, I-TS/TT appeared less broadly inflammatory than TF/TI but showed selective Th17-associated activity relative to TS/TT.

### The inflammatory response in active trachoma masks *Ct-*associated immune signals

Using Spearman’s rank correlation test, we independently examined whether *Ct* infection versus no infection was significantly associated with immune analytes independently within each grade (**Table 3**). A positive rho indicates that, within a given trachoma grade, *Ct*-positive samples tended to have higher analyte ranks than *Ct*-negative samples. The association between *Ct* infection and analyte concentration varied by grade, implying that the effect of infection may depend on the underlying immunological state of the tissue. TSLP was positively associated with *Ct* infection in all grades except I-TS/TT, while IL-13 and IFN-γ were positively associated in T0 and TS/TT, consistent with the ANCOVA results. The effect of *Ct* infection on the immune response could be discerned for the T0 and TS/TT grades but could not for TF/TI or I-TS/TT. These latter results are likely explained by the overall proinflammatory response in TF/TI and I-TS/TT that masks any defining response to *Ct*.

**Table 3.** Association of immune analytes with *Ct* infection compared to no infection independently for each trachoma grade using Spearman’s rank correlation test.

|  | T0 |  | TF/TI |  | I-TS/TT |  | TS/TT |  |
| --- | --- | --- | --- | --- | --- | --- | --- | --- |
| Analytes<sup>b</sup> | Spearman's $\rho$ | FDR | Spearman's $\rho$ | FDR | Spearman's $\rho$ | FDR | Spearman's $\rho$ | FDR |
| Eotaxin | 0.25 | 0.057 | -<sup>a</sup> | - | - | - | - | - |
| IFN-γ | 0.28 | **0.032** | - | - | - | - | 0.15 | **0.0098** |
| IL-10 | 0.31 | **0.018** | - | - | - | - | 0.14 | **0.011** |
| IL-13 | 0.36 | **0.0049** | - | - | - | - | 0.17 | **0.0019** |
| IL-17B | - | - | - | - | - | - | 0.15 | **0.0094** |
| IL-1α | 0.24 | 0.066 | - | - | - | - | - | - |
| IL-21 | 0.29 | **0.024** | - | - | - | - | - | - |
| IL-5 | - | - | - | - | 0.48 | 0.093 | 0.17 | **0.0019** |
| MCP-4 | - | - | - | - | - | - | -0.11 | 0.093 |
| MIP-1α | - | - | - | - | - | - | -0.18 | **0.0019** |
| MIP-1β | 0.24 | 0.07 | - | - | - | - | - | - |
| TNF-α | 0.3 | **0.023** | - | - | - | - | 0.14 | **0.011** |
| TNF-β | 0.31 | **0.018** | - | - | - | - | - | - |
| TSLP | 0.44 | **0.00025** | 0.5 | **0.0028** | - | - | 0.27 | **6.20E-08** |

<sup>a</sup> indicates an FDR >0.10
<sup>b</sup> Includes all analytes with FDR <0.10.

### Bernoulli mixture modeling (BMM) identifies four clinically distinct clusters that define unique Immune State Types (ISTs) among the population

To identify underlying immune profiles, we compared BMMs ranging from 2 to 8 clusters. The 4-cluster solution minimized the Bayesian information criterion (BIC = 18076.89), which evaluates goodness of fit for a statistical model while penalizing it for complexity, outperforming the 5-cluster (BIC = 18096.07) and 3-cluster (BIC = 18136.61) alternatives (**Fig 3A**). Based on this global minimum, we retained four clusters for subsequent profile characterization with sizes of n=162, 222, 129, and 154, for clusters 1–4, respectively.

**Fig 3.**
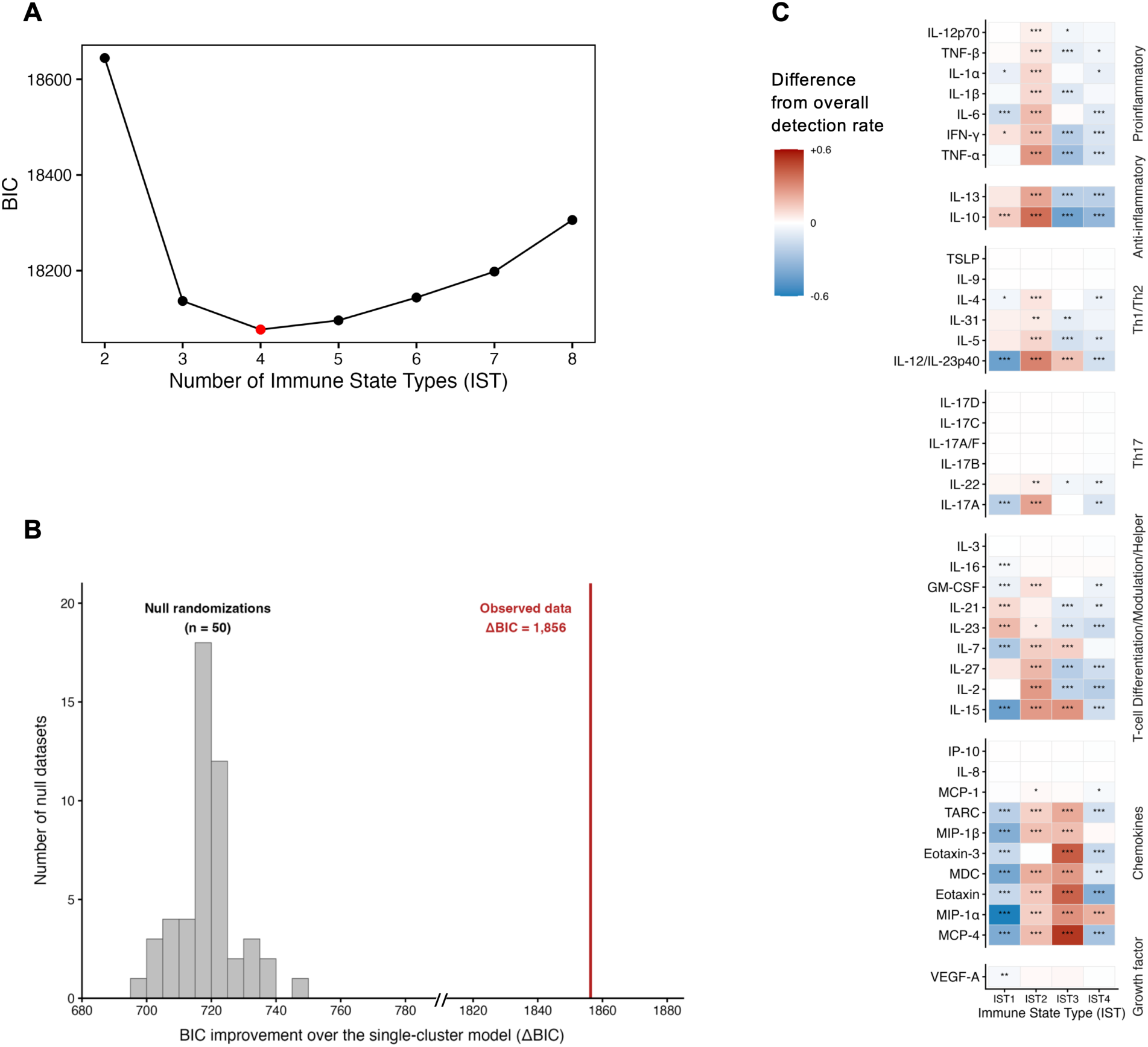
Immune State Type-specific analyte detection profiles. **(A)** Bayesian information criterion (BIC) values for Bernoulli mixture models (BMM) with 2 to 8 clusters, with the lowest BIC supporting the 4-cluster (IST) model. **(B)** Randomization test of IST structure, with istribution of ΔBIC from 50 null datasets compared with the observed ΔBIC. The observed improvement (1,856) greatly exceeded all null values. **(C)** Heatmap showing IST-specific differences in analyte detection rates relative to overall detection rates across all samples. Red indicates higher detection within an IST, blue indicates lower detection, and white indicates little or no difference. Analytes are grouped by functional immune category and ordered by variability across ISTs. Asterisks indicate significant IST-analyte differences after BH FDR correction: ***p<0.001, **p<0.01, and *p<0.05. Chi-square tests were used when all expected cell counts were ≥5; Fisher’s exact tests were used when any expected cell count was <5.

The observed improvement over a single-cluster model was then compared with 50 randomized matrices that preserved analyte prevalence and sample detection breadth. The observed improvement (ΔBIC =1,856) exceeded all null values (median 719, maximum 745; empirical *P* = 0.020) (**Fig 3B**), and no null matrix selected four clusters as optimal. This shows that the observed clustering reflects nonrandom patterns of analyte co-detection beyond those expected from analyte prevalence and sample detection breadth alone.

We tried five alternative clustering methods but none produced a partition that was both more stable and more clinically interpretable (see Methods). In particular, hierarchical clustering based on compositional distance did not recover the IST structure (adjusted Rand index, 0.06– 0.11). We therefore retained the Bernoulli mixture for downstream analyses.

The heatmap showed that the four ISTs differed in the relative detection of immune mediators (**Fig 3C**). IST2 showed an overall significant detection in most analytes relative to the detection rates across all samples. IST3 showed high detection of most chemokines, whereas IST1 and 4 showed more selective combinations of increased and decreased analyte detection, indicating that each IST represented a distinct immune profile.

Detection breadth differed across ISTs (P < 2 × 10⁻¹⁶), with a median of 21, 28, 25, and 21 analytes detected in IST1, IST2, IST3, and IST4, respectively. Recovered IST labels with only 54% cross-validated accuracy versus 92% for the full detection profile indicate that both breadth and analyte identity contributed to IST separation. Similar breadth in IST1 and IST4 (*P* = 0.68) further supported analyte identity as their primary distinction.

### Four unique ISTs define two scarring-associated and two inflammation/infection-associated immune profiles

The four ISTs comprised two inflammation/infection-associated profiles (IST1 and IST2) and two chronic trachoma-associated profiles (IST3 and IST4), each with distinct clinical and demographic characteristics. The distributions of trachoma grade, current *Ct* infection status, sex, and age are shown in **Fig 4** for each IST; enrichment and depletion results for trachoma grade and *Ct* infection are summarized in **Table 4**. Age differed significantly across the four clusters (Kruskal-Wallis test, *P* = 0.002), while sex did not.

**Fig 4.**
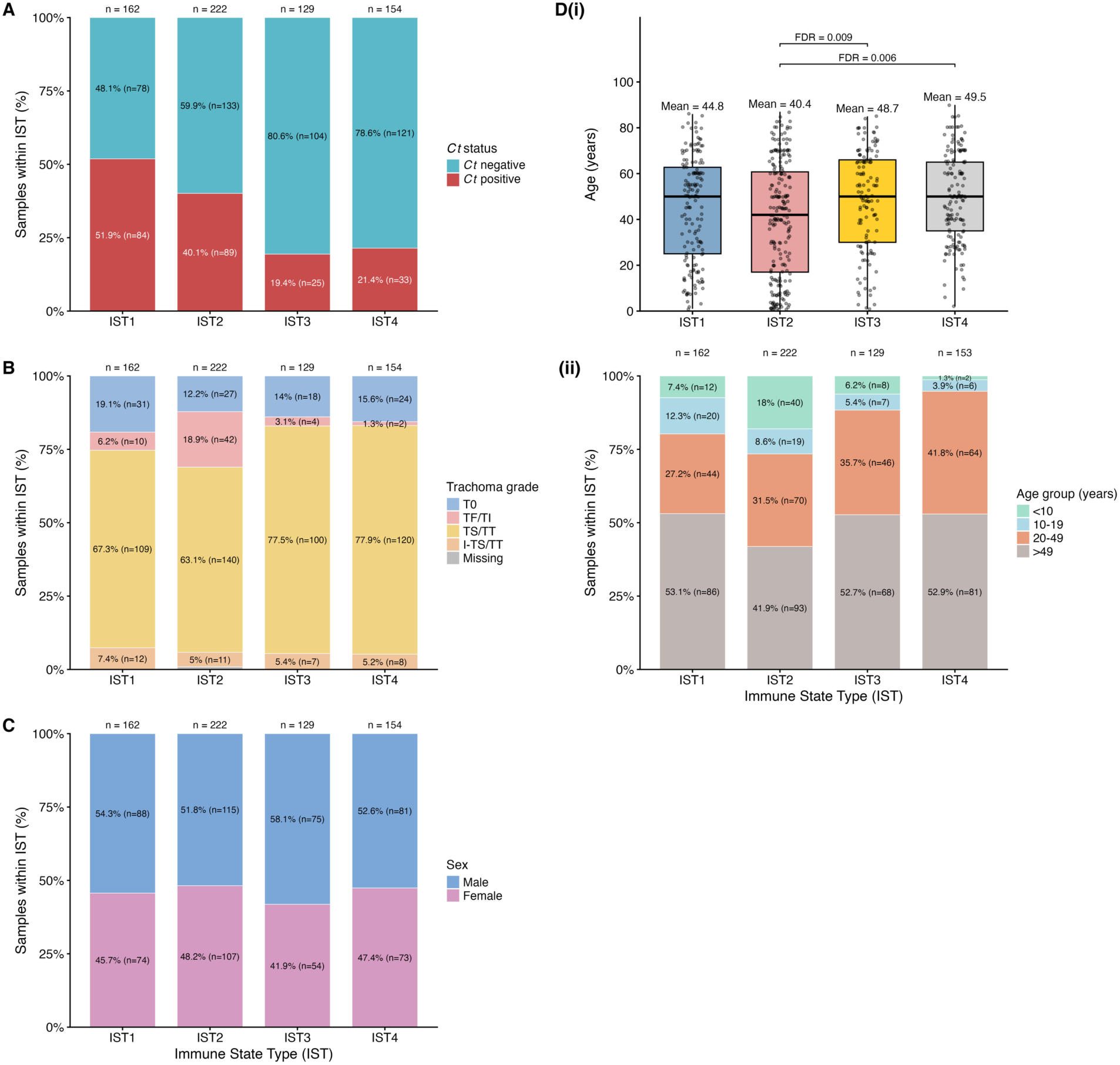
Clinical and demographic composition of Immune State Types (ISTs). Stacked bar plots show the distribution of **(A)** infection status, **(B)** trachoma grade, and **(C)** sex across ISTs. Values inside bars indicate the number and percentage of samples within each IST. **(D)** Age distributions across ISTs. (i) Participant ages by IST, with mean ages and significant FDR-adjusted pairwise Wilcoxon comparisons shown. (ii) Age-group composition within each IST; labels show percentages and sample counts.

**Table 4.** Enrichment and depletion of trachoma grades and *Ct* infection status across Immune State Types (ISTs).

| Tested category | IST1 (n=162) | IST2 (n=222) | IST3 (n=129) | IST4 (n=154) |
| --- | --- | --- | --- | --- |
| **Trachoma Grade** |  |  |  |  |
| **T0** | NS | NS | NS | NS |
| **TF/TI** | NS | 42/222 (18.9%); RR 5.26 (3.03-9.15), q=7.26e-10; Enriched | 4/129 (3.1%); RR 0.31 (0.11-0.84), q=0.0344; Depleted | 2/154 (1.3%); RR 0.12 (0.03-0.48), q=0.000814; Depleted |
| **TS/TT** | NS | 140/222 (63.1%); RR 0.85 (0.76-0.96), q=0.0126; Depleted | 100/129 (77.5%); RR 1.13 (1.01-1.26), q=0.0922; NS | 120/154 (77.9%); RR 1.15 (1.03-1.27), q=0.0462; Enriched |
| **I-TS/TT** | NS | NS | NS | NS |
| **Infection Status** |  |  |  |  |
| ***Ct*-** | 78/162 (48.1%); RR 0.68 (0.57-0.8), q=1.2e-6; Depleted | 133/222 (59.9%); RR 0.88 (0.78-1), q=0.0809; NS | 104/129 (80.6%); RR 1.31 (1.17-1.45), q=0.000336; Enriched | 121/154 (78.6%); RR 1.28 (1.15-1.42), q=0.00043; Enriched |
| ***Ct*+** | 84/162 (51.9%); RR 1.78 (1.46-2.18), q=1.2e-6; Enriched | 89/222 (40.1%); RR 1.26 (1.02-1.55), q=0.0809; NS | 25/129 (19.4%); RR 0.51 (0.35-0.73), q=0.000336; Depleted | 33/154 (21.4%); RR 0.56 (0.4-0.77), q=0.00043; Depleted |

IST1 and IST2 were associated with *Ct* infection and active trachoma and had younger age profiles, with mean ages of 44.8 and 40.4 years, respectively. Children under 10 years accounted for 7.4% of IST1 and 18.0% of IST2, compared with 9.3% of the overall dataset, indicating an enrichment of children in IST2 along with significant enrichment for TF/TI, which accounted for 18.9% of this IST; 42 (72.4%) of the 58 participants with TF/TI were assigned to IST2. IST2 also showed borderline enrichment for *Ct* positivity (40.1%, adjusted *P* = 0.08), consistent with an inflammatory, infection-associated state. IST1 had the highest *Ct* positivity (51.9%), the second-highest proportion of TF/TI (6.2%), and the highest proportion of I-TS/TT (7.4%). In addition, the two ISTs had substantially different immune profiles (**Fig 3C**). IST2 showed broadly higher detection across all analyte categories, whereas IST1 had a more quiescent profile characterized by higher detection of select cytokines, including IFN-γ, IL-10, IL-21, and IL-23, but significantly lower detection of all chemokines and other immune mediators. This is consistent with higher IFN-γ observed in *Ct-*positive samples in ANCOVA. Based on these profiles, we designated IST1 as “Quiescent Infection” and IST2 as “Broad Inflammation.”

IST3 and IST4 were distinct profiles associated with *Ct* negativity and chronic scarring. Participants in these ISTs were the oldest on average (48.7 and 49.5 years, respectively) with only 6.2% and 1.3% of children <10 years old, respectively. TS/TT and I-TS/TT accounted for and 82.9% and 83.1%, respectively, of each group compared with 76.0% of the overall cohort. IST4 was significantly enriched for TS/TT, while IST3 was borderline enriched (adjusted P = 0.09). Both ISTs were also significantly depleted of *Ct* infection and TF/TI. Immunologically, they shared a higher detection of the chemokines MIP-1α (**Fig 3C**), consistent with the elevated MIP-1α concentrations observed in TS/TT in the ANCOVA analysis. However, IST3 showed much more pronounced detection of chemokine activity than IST4, along with higher detection of several T-cell-associated cytokines, including IL-12/IL-23p40, IL-7, and IL-15, consistent with the elevated IL-7, IL-15, and chemokines in TS/TT in the ANCOVA analysis. We therefore designated IST3 as “Conjunctival Remodeling” and IST4 as “Quiescent Scarring,” representing two distinct scarring-associated states differentiated by the degree of ongoing immune activity.

### Geographic, but not occupational, factors align with IST composition

To examine whether environmental and occupational factors are associated with the observed immune heterogeneity, we tested whether these were enriched or depleted across the ISTs (**Fig 5**). IST composition did not differ significantly by adult occupation, and no adult occupational category was enriched or depleted. In contrast, individuals from village region 1 were depleted in IST3 (RR = 0.567, 95% CI: 0.411 to 0.783; FDR = 0.00148). Children, who were included in the occupation variable but represent a demographic rather than occupational group, were enriched in IST2 (RR = 4.81, 95% CI: 2.33 to 7.93; FDR = 1.19 × 10⁻⁵) and depleted in IST4 (RR = 0.0775, 95% CI: 0.0108 to 0.558; FDR = 0.0111). This pattern is consistent with the age distributions described above and with prior findings that children have a higher risk of *Ct* infection [28]. Together, these results indicate that ISTs differ by demographic and geographic composition, in addition to trachoma grade and infection status, but not by adult occupation.

**Fig 5.**
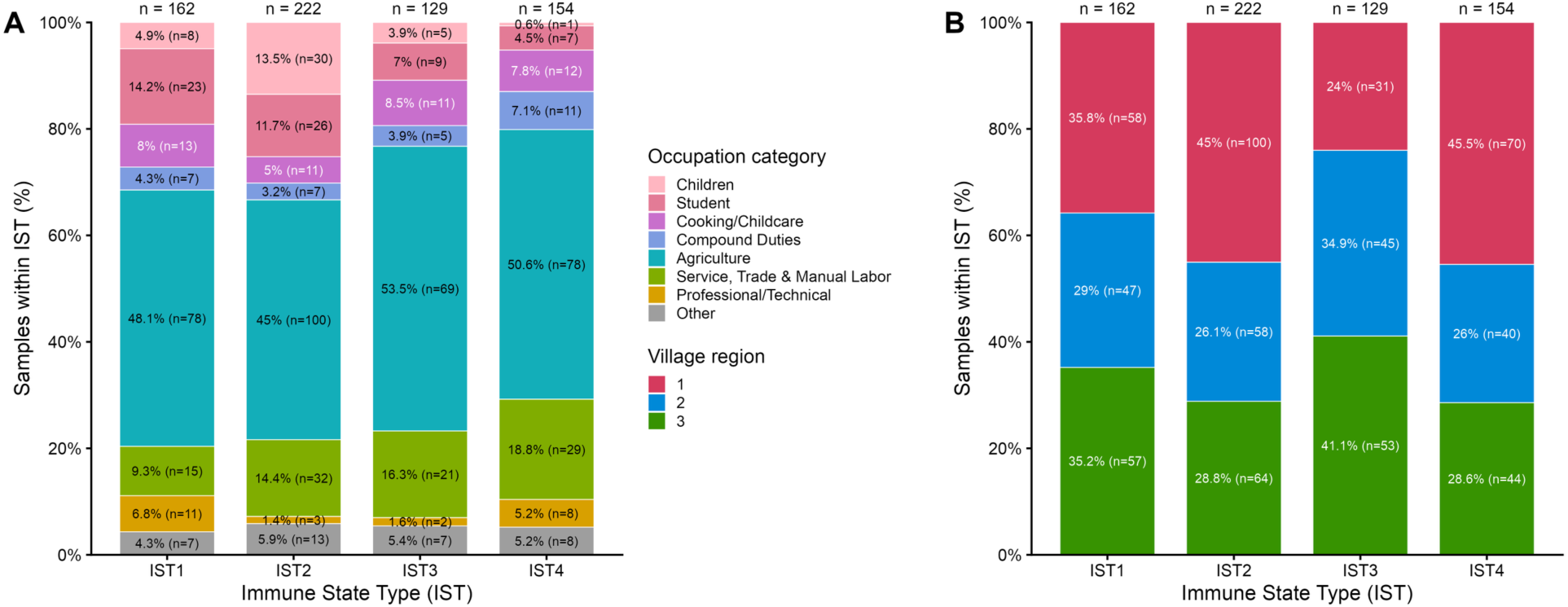
Occupation category and village region composition by IST. Stacked bar plots show the percent composition of each IST by **(A)** occupation category and **(B)** village region. Labels indicate the sample count and percentage within each cluster.

## DISCUSSION

Trachoma pathogenesis does not follow a simple linear sequence in which *Ct* infection induces inflammation and repeat infection ultimately leads to scarring [2]. In the hyperendemic population of Amhara, quantitative protein profiling revealed that *Ct* infection, active trachoma, chronic scarring with and without inflammation were each governed by distinct immune response profiles mirrored in four unique ISTs. While the same modeling tool has resolved complex subphenotypes in conditions like long COVID [47], and plasma cytokine profiling based on other clustering methods identified disease-associated immune states in asthma [48] and COVID-19 [49], this is the first study to define conjunctival ISTs. Here, the ISTs captured biological variation that could ultimately identify individuals before a full-blown immune response ensues (IST1) and those with profiles potentially relevant to pathogenic disease progression to TT (IST3) for a novel immune or fibrotic therapeutic or other intervention.

Current *Ct* infection was associated with Th1 cytokine inflammation (IFN-γ) and a robust Th2-associated mucosal repair response (TSLP, IL-13, IL-9, and IL-17B). Spearman rank correlations supported the ANCOVA findings while revealing some variation: TSLP was significantly higher in *Ct*-positive samples for all grades except I-TS/TT, while IFN-γ, IL-13 and IL-17B were higher in T0 and in TS/TT. However, there were fewer analytes associated with infection for TF/TI or for I-TS/TT. These findings indicate that pronounced inflammation, particularly in TF/TI, likely obscures or dampens any further immune response to *Ct* infection.

The elevated IFN-γ in *Ct* infection agrees with previous conjunctival mRNA expression studies [4–11] and its role in controlling intracellular bacteria like *Ct* [50]. The type 2 profile is consistent with an alarmin–driven Th2 repair pathway initiated by TSLP release following epithelial damage, which promotes IL-13-producing Th2 responses [51]; IL-13 limits inflammatory injury and supports epithelial and extracellular matrix repair [52]; Th2-derived IL-9 can amplify type 2 responses [53]; and IL-17B, a lesser-known IL-17 family cytokine, can stimulate additional Th2 cytokine production [54]. This is consistent with previous reports of increased mRNA expression of the Th2 cytokine IL-4 in infected conjunctiva [5]. However, repeated activation of the same repair program may promote collagen deposition and fibrosis, as observed in schistosomiasis and in stronger type 2 responses to *Ct* antigens among individuals with scarring trachoma [55,56]. Thus, the *Ct*-associated TSLP and IL-13 response may be protective during acute injury but could become maladaptive if repeatedly activated. Despite these selective Th1– and type 2-associated signals, *Ct* infection did not involve generalized inflammatory activation. IST1, the most *Ct*-enriched state, showed elevated IFN-γ detection but little chemokine or broader immune mediator activity consistent with the ANCOVA analyses and was therefore designated “Quiescent Infection.”

TF/TI revealed broad proinflammatory and Th1/Th17 activity with early remodeling. Higher IFN-γ, TNF-α, TNF-β, MIP-1α, and MIP-1β supported Th1 activation [57–59], consistent with cell-mediated control of active trachoma [60]. IFN-γ can induce the depletion of tryptophan, an essential amino acid for *Ct* development and growth, driving the organism into a persistent state and potentially prolonging inflammation [61,62]. Concurrent IL-17A and IL-22 upregulate support for Th17-associated inflammation, while IL-17B/C/D indicate broader IL-17 axis modulation [63]. IL-17A, active at epithelial barriers and previously observed to be elevated in active trachoma, correlated with profibrotic connective tissue growth factor (CTGF) [64]; IL-22 may limit IL-17-mediated epithelial damage, as observed in experimental dry eye disease [65]. Overall, these responses support antimicrobial defense while promoting ocular inflammation and early remodeling.

The concurrent elevation of IL-17A, IFN-γ, IL-12/IL-23p40, and GM-CSF also suggests Th17-to-Th1 plasticity. IL-12 and IL-23 share the p40 subunit and promote Th1 and Th17 differentiation, respectively [66]. IL-12 can drive Th17-to-Th1 transition through an IL-17A+/IFN-γ+ intermediate [67]. These Th1-like cells are more pathogenic than conventional Th17 cells as described in rheumatoid arthritis and also occur in Crohn’s disease, multiple sclerosis, and type 1 diabetes [68–71]. They may contribute to chronic conjunctival inflammation, although cell-resolved studies are needed to establish their presence.

Elevated IL-15, IL-16, and IL-10 in TF/TI suggest coordinated T-cell regulation. IL-16 recruits and activates CD4+ T cells [72] while IL-15 sustains CD8^+^ and CD4^+^ T cells [73], and IL-10 suppresses antigen-presenting cell activity to dampen Th1 inflammation [74]. This agrees with previous reports of increased IL-10 in active trachoma with *Ct* and persistent expression of regulatory T-cell function-related transcription factor forkhead box P3 (*FOXP3*) after *Ct* clearance [8].

Broad chemokine elevation in TF/TI further suggests immune recruitment and remodeling. Eotaxins, MDC, MCP-1, MIP-1α, MIP-1β, and IL-8 are linked to fibroblast migration, activation, or fibrosis in other tissues [75–80]. MCP-1 participates in the fibrotic process from fibrosis initiation through progression 9/30/26 2:37:00 PM, while IL-8 may similarly activate conjunctival fibroblasts as it does dermal fibroblasts in pathological scarring following commensal dysbiosis [80]. Higher VEGF-A may reinforce this environment. Based on immunohistochemistry, it is upregulated in active trachoma [81] and promotes transforming growth factor (TGF)-β1 signaling and myofibroblast transformation in experimental rabbit subconjunctival scarring, while VEGF inhibition reduces fibrosis [82]. Increased vascular permeability may further facilitate immune-cell infiltration [83]. Overall, TF/TI reflects broad mucosal immune activation, combining Th1/Th17 signaling, regulatory counter-responses, and remodeling that may precede scarring, consistent with the increased expression of scarring-related pathways, such as metalloproteinase activity and collagen formation, during active trachoma [5,7]. IST2 recapitulated this profile, showing strikingly high detection across immune analytes for most TF/TI participants.

Far less is known about inflammation in established scarring, despite longitudinal studies in Ethiopia and Tanzania linking recurrent inflammatory episodes to scarring progression [21]. Females had a much higher risk of I-TS/TT compared to males, consistent with their higher trachoma burden in endemic regions and repeated exposure to infected children through caregiving [84]. Yet, only two previous studies directly examined conjunctival immunology in this context: A 1996 study of I-TS/TT did not provide any insight into this grade and was limited by a sample size of seven and narrow cytokine transcript panel [6]. Our 2008 25-analyte protein study found higher IL-1β and lower Interleukin-1 receptor antagonist (IL-1Ra) and IL-12p40 levels in TT/TI compared with T0 [17]. While we found IL-17-associated inflammation, T-cell modulation, chemokine recruitment, and VEGF-A-associated remodeling, the Th1/Th17 responses were attenuated relative to TF/TI, consistent with our previous finding of reduced Th1 immunity in TT/TI [17]. Compared with TS/TT, I-TS/TT had higher IL-17A and lower IP-10, further suggesting selective Th17 reactivation without a broadly increased IFN-γ-induced chemokine signaling [85]. Thus, I-TS/TT may represent a Th17-skewed inflammatory state within an established scarring environment. I-TS/TT distribution varied little across ISTs, possibly due to the smaller sample size of 38. Larger studies of this intermediate clinical state are needed to determine whether and where it sits within in a distinct IST.

TS/TT showed a narrower but persistent injury and remodeling profile, consistent with biopsy evidence of greater inflammatory-cell infiltration in scarred than control conjunctiva [9,86], confirming that scarred tissue is far from immunologically silent. Elevated T-cell modulatory cytokines IL-7, IL-15, and IL-16 suggest continued T-cell maintenance and recruitment that may sustain chronic inflammation without clinically active disease. Elevated IL-1α, an alarmin released after cellular damage [82], in TS/TT may link ongoing epithelial injury to remodeling through its capacity to induce proinflammatory fibroblasts and collagen accumulation as in lung injury [87]. IL-5 may promote remodeling through eosinophil activation, enhancing fibroblast proliferation, collagen synthesis, and lattice contraction partly via TGF-β [88]. Reduced eosinophilic inflammation and airway fibrosis in IL-5-deficient mice supports a potentially similar role in the scarred conjunctiva [89].

The chemokine profile in TS/TT further supports persistent remodeling. Elevated IL-8, MCP-1, MIP-1α, MIP-1β, eotaxin, and eotaxin-3 and similar increases in all but eotaxin-3 were observed in our previous study [17]. These chemokines have been implicated in fibrosis-related biology in other tissues [54,75,78,79,90]. Together with IL-1α and IL-5, they suggest that immune-cell recruitment and fibrogenic activity continue after scars become clinically visible. Furthermore, compared with TF/TI, TS/TT showed broadly attenuated proinflammatory, Th17, T-cell, regulatory, and chemokine signals while selected injury and remodeling pathways persisted. Here, scarring appears to develop not through further amplification of inflammation, but through continued remodeling after inflammatory activity has waned.

The heterogeneity within TS/TT resolved into IST3 (Conjunctival Remodeling) and IST4 (Quiescent Scarring). IST3 showed stronger chemokine and T-cell-associated activity, including higher IL-7, IL-15, IL12/IL23p40, suggesting both ongoing remodeling and maintenance of an inflammatory tissue environment. In contrast, IST4 represents a quiescent or “exhausted” state in which MIP-1α persists but broader chemokine and T-cell-associated activity has declined. This difference could potentially help explain the wide variation in scar burden within TS/TT. A recent photographic grading study showed that conjunctival scarring can range from short lines of scars to scarring involving ≥90% of the conjunctiva [91]. Together, the four ISTs provide a conceptual framework for the distinct immune profiles associated with *Ct* infection, active inflammation, and scarring with and without remodeling (**Fig. 6**).

**Fig 6.**
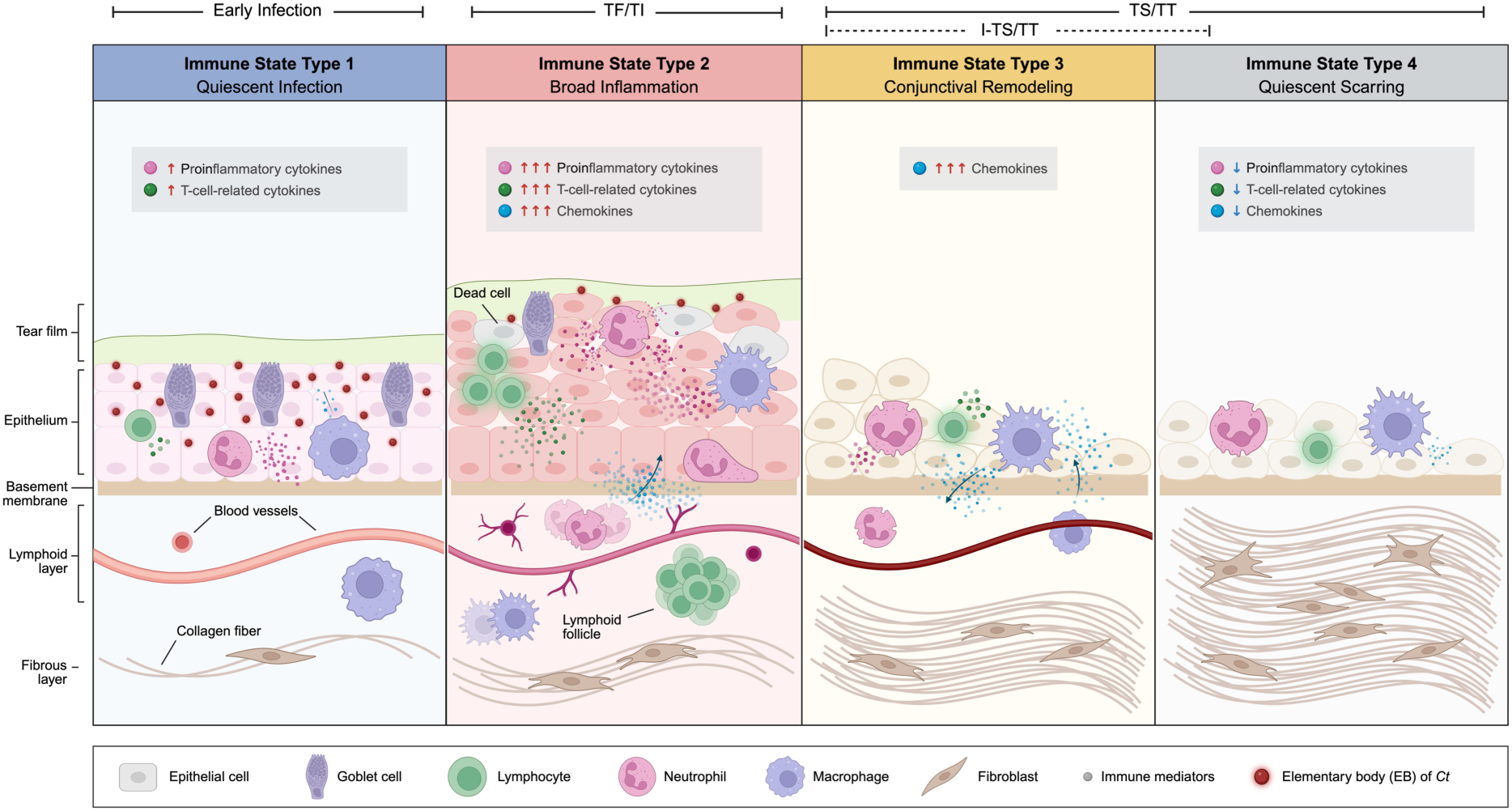
Conceptual model of conjunctival ISTs across *Ct* infection, inflammation, and scarring. Four panels pair BMM-defined ISTs with representative histologic features reported for the corresponding clinical states [92]. IST1, “Quiescent Infection,” shows selective detection of proinflammatory and T-cell-related cytokine with relatively preserved epithelium, goblet cells, and basement membrane. IST2, “Broad Inflammation,” shows widespread cytokine and chemokine detection, epithelial hyperplasia and injury, increased immune-cell infiltration, and lymphoid follicle formation. IST3, “Conjunctival Remodeling,” shows persistent chemokine activity, epithelial and goblet-cell loss, reduced tear-film support, immune-cell recruitment, fibroblast activity, and collagen accumulation. IST4, “Quiescent Scarring,” shows reduced immune activity and established fibrosis. Clinical brackets indicate the nonexclusive associations between the ISTs and early infection and trachoma grades. The small I-TS/TT sample (n=38) limits its precise placement within the IST framework, although its inflammatory, scar-associated profile conceptually aligns most closely with IST3. Created with BioRender.com.

In conclusion, protein-level mapping identified immune profiles associated with *Ct* infection, inflammation, and scarring while also resolving clinical disease into relevant ISTs. With further validation and reduction to a practical biomarker panel, IST1 could identify a infection-dominant phase in which antibiotic treatment should remain central, whereas the widespread host immune response in IST2 supports the investigation of adjunct immunomodulatory therapy to limit inflammatory damage and disease progression. The chemokine profiles separating IST3 from IST4 may support a biomarker panel that distinguishes active from quiescent scarring and, with validation, help identify individuals who may be at risk of progression to TT and benefit from novel antifibrotic intervention. As global programs pursue the WHO target of eliminating trachoma as a public health problem by 2030, this approach may become especially valuable as efforts shift from population-level control to identifying residual infection and individuals at continued risk of vision loss. Longitudinal and cell-level studies are needed to validate the prognostic value of the ISTs and identify modifiable host pathways that can be therapeutically targeted before vision-threatening pathology ensues.

## ACKNOWLEDGEMENTS

We are grateful to the Amhara community members who contributed to this research and to the Bahir Dar Specialty Eye Center staff for supporting field studies and advancing eye care in the region. Funding was provided by the National Institutes of Allergy and Infectious Diseases grant R01AI158527 (to DD and TDR). The funder was not involved in developing or conducting the study, evaluating or interpreting the data, or deciding to publish the findings.

## AUTHOR CONTRIBUTIONS

X.C.S.: Investigation, Methodology, Software, formal analysis, validation, visualization, writing—original manuscript, review and editing.

O.O.: Investigation, Methodology, Software, Analysis, Validation, Writing – review & editing

B.R.T.: Formal analysis, Investigation

S.C.: Methodology, Validation

H.W.D.: Investigation, methodology, supervision

K.A.: Investigation, methodology, supervision

T.D.R.: Conceptualization, Funding acquisition, Investigation, Resources, Writing – review & editing

D.D.: Conceptualization, Funding acquisition, Investigation, Methodology, Visualization, Project administration, Resources, Supervision, Writing – review & editing

## SUPPORTING INFORMATION CAPTIONS

**Table S1. Participant metadata, IST assignments, and conjunctival immune analyte concentrations**.

## Notes

### Competing Interest Statement

The authors have declared no competing interest.

